# Isolation and characterization of microbiota from human pancreatic tumors and small intestine

**DOI:** 10.1101/2024.11.27.625762

**Authors:** Dominik Awad, Holly Attebury, Ryan Hong, Kwi Kim, Li Zhang, Allison Bischoff, Aaron deDekker, Matthew Hoostal, Joseph A. Nieto Carrion, Noah S. Nelson, Chris Strayhorn, Timothy Frankel, Marina Pasca di Magliano, Costas A. Lyssiotis, Thomas M. Schmidt, Donnele Daley

## Abstract

Pancreatic ductal adenocarcinoma has a unique tumor microbiome and the systemic depletion of bacteria or fungi using antibiotic/antifungal cocktails leads to a decrease in pancreatic tumor burden in mice. However, functional studies remain rare due to the limited availability of clinically relevant microbiota. Here, we describe in detail the isolation of bacteria and fungi from the small intestine and tumor of pancreatic cancer patients at the Rogel Cancer Center. We then further characterized the impact of a newly isolated *Klebsiella oxytoca* strain (*UMKO1*) on the pancreatic tumor microenvironment using bacterial genome sequencing, untargeted and targeted metabolomics, as well as an *ex vivo* tumor transplant system. We found that *UMKO1* possesses a gene for the long form of cytidine deaminase, which can inactivate the standard PDAC chemotherapeutic agent gemcitabine. In addition, we found that *UMKO1* can produce several indoles when grown in tumor-like conditions, metabolites that can lead to an immune suppressive environment and interfere with therapy outcome. To test this in detail, we assessed changes in immune populations in pancreatic tumor explants upon exposure to the supernatant of *UMKO1* and other isolated bacteria grown in tumor Interstitial fluid media (TIFM). We found that while none of the bacterial supernatants changed the abundance of CD8 T cells, granzyme B positive CD8 T cells were the lowest in tumor explants exposed to *UMKO1*, and not other isolated *Klebsiella* species or the non-pathogenic laboratory strain *E. coli K12*. In summary, the isolated collection of bacteria and fungi from this study are a valuable toolbox to study the impact of microbiota on pancreatic cancer.

## Introduction

Pancreatic ductal adenocarcinoma (PDAC) is one of the deadliest major cancers with an overall 5-year survival rate of just 13%^1^. This is in part due to the lack of biomarkers for early detection, but also because of its resistance to chemo- and immunotherapy. PDAC is surrounded by a dense stroma, mainly consisting of cancer-associated fibroblasts and immunosuppressive myeloid populations. In addition, PDAC has its own unique microbiome^2–8^. Interestingly, long-term survivors of pancreatic cancer demonstrate increased diversity in their tumor microbiome, as well as an enrichment of several bacteria such as *Bacillus*, *Pseudoxanthomonas, Saccharopolyspora* and *Streptomyces*^5^. Ablation of the gut and tumor microbiome in mice via antibiotic and antifungal cocktails decreased pancreatic tumor burden in multiple studies and this phenotype could be reversed with murine or human fecal microbial transplants (FMT)^3–6,8^. In particular, FMT from short term survivors or from mice harboring a pancreatic tumor increased tumor growth in mice, while FMT of long term survivors restricted tumor growth^3,6^. While the studies have revealed important insights into the role of the gut and tumor microbiome, studying the impact of individual bacteria or fungi remains a limitation in our understanding. The low biomass of microbes in the tumor do not allow for reliable metagenomic sequencing, therefore limiting the information to general analysis and providing little information about their genome. To overcome this challenge and study pancreatic cancer-associated microbiota in more detail, we isolated bacteria and fungi from the small intestine and the pancreatic tumor of pancreatic cancer patients at the University of Michigan Rogel and Blondy Pancreatic Cancer Center undergoing surgical resection. We demonstrated herein that we can cultivate various bacteria and fungi species from both tissues, which we have made available to the research community. Further, we characterized the activity and functions of the most abundant isolated bacteria in the pancreatic tumor, *Klebsiella oxytoca,* using bacterial genomic sequencing, untargeted and targeted metabolomics, as well as an optimized ex vivo tumor explant system. By creating a broad collection of clinically relevant microbiota isolated from human pancreatic cancer patients, we are able to provide new resources to study the impact of individual bacteria and fungi on pancreatic tumors.

## Materials and methods

### Tissue collection

#### Tumor and small intestine from pancreatic cancer patients

Pancreatic tumor and duodenum tissue were collected after written patient consent under the approved IRB protocol (HUM00025339, PI:Vaibhav Sahai, Co-I: Dr. Donnele Daley). The tissue was either flash frozen or frozen in cryotubes containing RPMI media and stored at -80 °C until the start of the isolation. Additional tissue samples were flash frozen for 16S/ITS sequencing.

#### Donor pancreata

Donor pancreata and small intestines used for this study were not eligible for organ donation or did not have a recipient and were collected from the Gift of Life Michigan Donor Care Center. The tissue was processed as previously described in detail^9^. The use for scientific research was approved by the Gift of Life research review group (PIs: Dr. Timothy Frankel, Dr. Marina Pasca DiMagliano). For the isolation we used the section of the duodenum (small intestine closest to the pancreas) and the tissue from the head of the pancreas.

### Tissue processing for microbial isolation

Tissue previously frozen in RPMI was thawed on ice and flash frozen tissue was thawed on ice after the addition of sterile RPMI. Tissue was then cut into equal size pieces in a Baker Class II Biosafety Cabinet designated for BSL2 microbes. After dissection, tissue pieces were added to individual media plates and streaked with sterile forceps before moving the plates to the hypoxic chamber. For anoxic conditions, the tissues were streaked in the anoxic chamber directly. Plates were incubated in Coy glove boxes (Coy Laboratory Products Inc., Grass Lake, MI) for isolation and cultivation of microbes at 37°C under anoxic (5% CO2, 1.5 to 3.5% H_2_ and N_2_ balance) and hypoxic (2% O_2_, 5% CO2, balance N2) conditions. Mixed culture plates were inspected for colony formation every 12 hours and colonies were restreaked to avoid overgrowth. Early optimization work of this isolation process did not show any benefit of plating or spotting the concentrated RPMI solution from tissue storage onto plates.

### Media composition

All media components below are listed as quantities needed to prepare 1 L of media. If not otherwise stated, dissolve into deionized water to reach a final volume of 1000 ml.

#### Sabouraud agar

Peptone 10 g (BD Cat #211693), Glucose 40 g (Acros Organics Cat #41095-5000), Agar 15 g (Invitrogen Cat #30391-023); bring with deionized water to 1000 ml and adjust pH to 5.6.

#### BHI Agar

BHI agar 52 g (BD Cat #211065) + Calcium Chloride 2.2 g (Sigma-Aldrich Cat #C8106-500G)

#### Bacillus clausii Agar

Peptone 2g (BD Cat #211693), Yeast extract 5 g (Fisher Scientific Cat #BP1422-500), Glucose 1 g (Acros Organics Cat #41095-5000), K_2_HPO_4_ 1 g (Sigma-Aldrich Cat #P8281-500G), MgSO_4_.7H_2_O 0.2g (Sigma Life Science Cat #63138-250G), Skim Milk 2 g (BD Cat #232100), Saline solution 0.9% (Sodium chloride, Sigma Aldrich Cat #S9625-500G), Ampicillin 50 mg (Fisher Bioreagents Cat #BP902-25), Nystatin 5 mg (Sigma-Aldrich Cat #N6261).

#### Modified Columbia Agar

Columbia Broth 35 g (Remel Cat #R452972), 5% Sheep’s blood (Remel Cat #R54012), 0.5% NaCl (Sigma Aldrich Cat #S9625-500G), Tryptophan 40 mg (Sigma Life Science Cat #T8941-100G), Histidine 40 mg (Sigma Cat #H5659-25G), Agar 15 g (Invitrogen Cat #30391-023)

#### Gifu Anaerobic Medium (GAM) Agar

GAM Broth 59 g (HiMedia Cat #M1801-500G), Agar 15 g (Invitrogen Cat #30391-023)

#### Leeming-Notman Agar

Bacteriological peptone 10 g (BD Cat #211693), Glucose 10g (Acros Organics Cat #41095-5000), Yeast extract 2 g (Fisher Scientific Cat #BP1422-500), Ox bile, desiccated 8g (Fluka Analytical Cat #70168-100G), Glycerol 10 ml (Fisher Scientific Cat #G33-500), Glycerol monostearate 0.5 g (Thermo Scientific Cat #043883.30), Olive Oil 20.0 ml (Kirkland Signature), Agar 15 g (Invitrogen Cat #30391-023) based on^9^. Adjust pH to 6.2 with 1M HCl. Add 5 ml of warmed Tween 40 (60-70C) (TCI Cat #T0544), boil briefly to dissolve components and then autoclave.

<u>CHROMagar™ Malassezia</u> was used as directed by the supplier.

### MALDI-TOF

Colonies of isolated bacteria and fungi were added directly to a 1.5 mL Eppendorf tube using a 1 µL loop along with 300 µL of UHPLC water. Afterward, 900 µL of absolute ethanol was added, and the colonies were centrifuged at 13500 rpm for 2 minutes. The supernatant was removed, and the pellet was dried on a heating block at 40°C for 15 minutes. Once dried, the pellet was resuspended in 25 µL of 70% formic acid and mixed by pipetting. Following this, 25 µL of acetonitrile was added, and again mixed by pipetting. The suspension was then centrifuged at 13,500 rpm for 2 minutes. A total of 1 µL of the supernatant was transferred to an MBT Biotarget 96 Chip (Bruker), air-dried, and overlaid with 1 µL of HCCA matrix solution. Once dried, the plate was transferred to the MALDI-TOF MS instrument (MALDI Biotyper Sirius) and analyzed using the MBT Compass and Flex Control software.

### Genome sequencing of *K. oxytoca*

Bacterial genome sequencing was performed by Plasmidsaurus using Oxford Nanopore Technology with custom analysis and annotation. FASTA files for other Klebsiella strains were downloaded from the National Library of Medicine Nucleotide database (https://www.ncbi.nlm.nih.gov/nuccore). FASTA files were annotated using Prokka^10^ and the Genbank annotation files were visualized using Pan-genome Explorer^11^.

### Metabolomics

#### Sample prep

*Klebsiella oxytoca (UMKO1)* was grown on the modified Columbia agar (see media composition) and prior to inoculating in Tumor interstitial fluid media (TIFM), washed three times with PBS to avoid any carry over of metabolites from the media. *UMKO1* was grown for 16h while shaking at ambient temperatures. Supernatant was then collected by spinning the cells for 5 min at 3000g and filtered through a 28mm syringe filter (Corning, 431229, 0.2 µM PES). TIFM media control was subjected to the same incubation, spin and filtering. 400 ul TIFM or bacterial supernatant was then crashed with ice cold methanol, resulting in 4:1 methanol:media mix. The samples were then incubated on dry ice for 10 minutes and spun down at 11.000 x g in a precooled centrifuge for 10 minutes. The supernatant was dried using a speedvac and reconstituted in 80 ul of methanol:water (1:1) and transferred to a 300 ul insert with polyspring (JSI Scientific) and the insert was transferred to a phenomenex vial (AR0-9921-13-C) and capped (AR0-8952-13-B). Controls included negative controls such as matrix blank, solvent blank. A pool of all samples was used for the QC and identification in the untargeted analysis.

#### Untargeted metabolomics

LCMS grade of acetonitrile (Supelco), methanol (Sigma), ammonia (Millipore), Isopropanol (Fisher), formic acid (Fisher), ammonia acetate (Sigma) were used for these experiments. The water was from a Millipore purified filter device with 18 omegas. Thermo Scientific IQX Orbitrap LC-MS system consists of two Vanquish Horizon UHPLC binary pumps, a dual Vanquish autosampler, two Vanquish column compartments with a switch valve for two column setup configurations and a Thermo Multiwavelength Detector CG and the IQX Orbitrap HRMS. Thermo Scientific Excalibur Version 4.6.67.17 data system with Dell computer Opex-EX3 with Winindo Professional 10 operating system was used for tuning, calibration, optimization and data acquisition. Data processing was performed by Thermo Scientific Tracing Finder 3.1, FreeStyle for initial analysis and untargeted data was then further analyzed using Compound Discover 3.3.1.111 using the *Untargeted Metabolomics workflow* with modifications. For untargeted metabolomics applications, the Waters Acquity UPLC BEH amide 1.7 um 2.1 x 100 mm column with a Waters UPLC BEH Amide 1.7 um VanGuard Pre-Column column of amide for high throughput acquisitions. Solvent A consists of 10 mM ammonium acetate in water with 10 mM Ammonia pH 9.2. Solvent B consists of acetonitrile. Pump Seal wash and autosampler wash is 50% isopropanol with 0.1% formic acid. The LC profile is 0.2 ml/min of 85% B at 0-0.5 min: 0.5-15 min, 15%B; 15-17 min, 15% B: 17.1 min, 85%B; 22 min, 85% B. The column compartment temperature is maintained at 30 °C and the autosampler is at 5 °C. The injection volume is 3 µl.Thermo Scientific IQX MS is calibrated with Pierce IQX Orbitrap MS is calibrated weekly by Pierce FlexMix Calibration Solution (A39329) to with 2 ppm for all calibrates with infusion into the micro ESI probe at 5 µl/min. Source parameters: H-ESI Positive Static Voltage 4000 V; Negative Static Voltage 3500 V; Sheath Gas (Arb) 40, Aux Gas 5 (Arb), Sweep Gas 1 (Arb); Ion transfer Tube Temp 300 °C and Vaporizer Temp 350 °C. Orbitrap resolution is 60K, mass scanning range is 50-1000 da. Data type is centroid and polarity swathing mode of acquisition is applied.

#### Targeted metabolomics

LCMS grade of acetonitrile (Supelco), methanol (Sigma), ammonia (Millipore), Isopropanol (Fisher), formic acid (Fisher), ammonia acetate (Sigma) were used for these experiments. The water was from a Millipore purified filter device with 18 omegas. Reference standards used in this study were kynurenic acid (TCI Cat# H0303), L-kynurenine (TCI Cat #K0016), indole-3-pyruvic acid (Sigma Cat# I7017), DL-indole-3-lactic acid (Sigma Cat#I5508), tryptophan and indole-3-acetic acid, and were acquired from Cayman Chemicals. Thermo Scientific Triple Quad TSQ Quantis LC/MS/MS system consists of a Vanquish binary pump, a Vanquish autosampler, and a Vanquish column compartment with a switch valve for two column setup configurations. Thermo Scientific Excalibur Version 4.4.16.14 data system with Dell computer Opex-EX3 with Win10 operating system was used for calibration, compound optimization and sample data acquisition. Data processing is performed by Thermo Scientific Trace finder 3.1 and Skyline 12.1. A Thermo Hypersil Gold aQ C18 2.5µm 2.1 X 100 mm Column and a Phenomenex High Pressure column protection filter is used for the separation. Solvent A consists of Water with 0.1% FA. Solvent B is pure acetonitrile. Pump Seal wash and autosampler wash is 50% isopropanol with 0.1% formic acid. The LC profile is at 0.15 ml/min, 0-1 min, 1%: 1 min, 90%B; 10min, 90% B: 12 min, 1%B; 12.1 min, 1% B; 15 min, 1% B. The column compartment temperature is maintained at 30 °C and the autosampler is at 5 °C. Injection volume is 3 µl. TSQ Quantis Triple Quad MS is calibrated with Pierce Triple Quadrupole Calibration Solution External Mass Range with microESI probe from Thermo Scientific. Infusion rate is at 3 µl/min. Source parameters: H-ESI Positive Static Voltage 3500 V; Sheath Gas (Arb) 40, Aux Gas 5, Sweep Gas 1; Ion transfer Tube Temp 325 °C and Vaporizer Temp 350 °C. Dewell Time is 80 ms. Q1 and Q3 FWHM are at 0.7 da. CID gas (mTorr) is at 1.5 with helium.

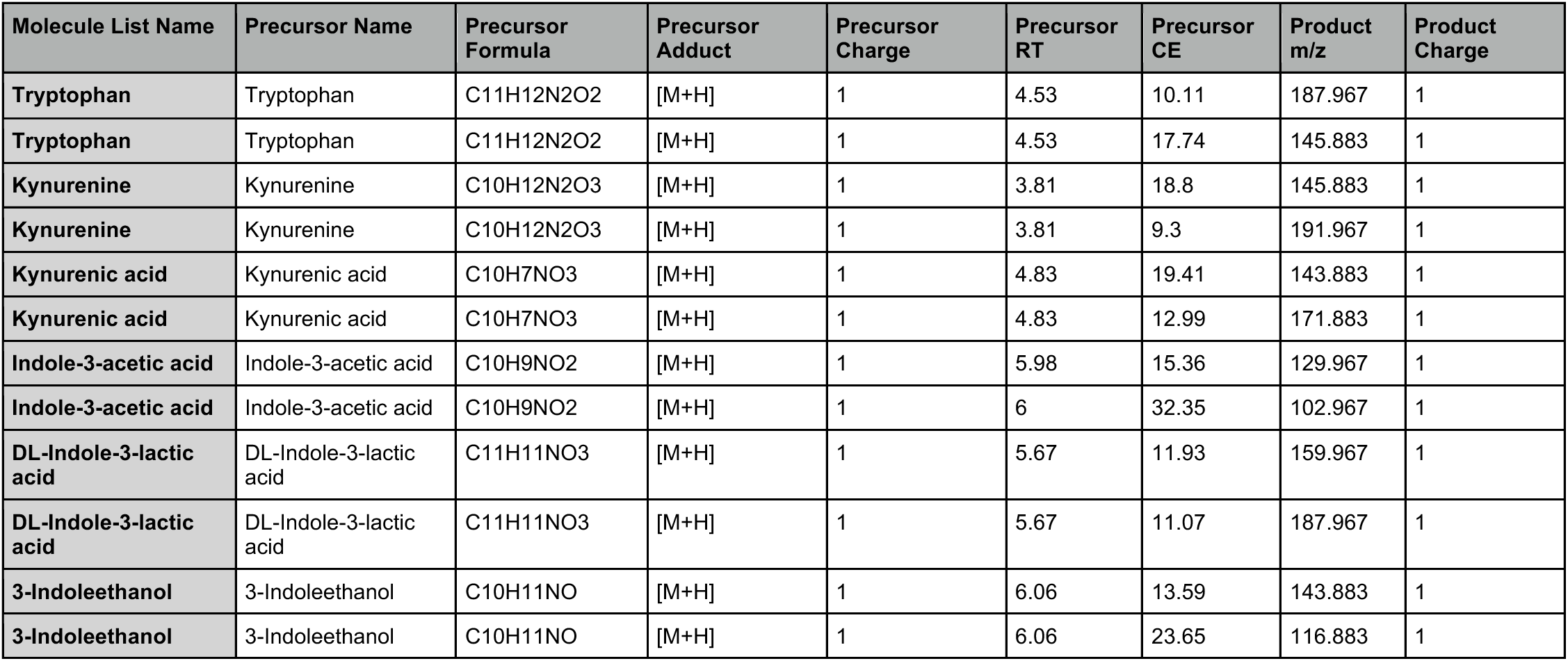

| Molecule List Name | Precursor Name | Precursor Formula | Precursor Adduct | Precursor Charge | Precursor RT | Precursor CE | Product m/z | Product Charge |
| --- | --- | --- | --- | --- | --- | --- | --- | --- |
| Tryptophan | Tryptophan | C11H12N2O2 | [M+H] | 1 | 4.53 | 10.11 | 187.967 | 1 |
| Tryptophan | Tryptophan | C11H12N2O2 | [M+H] | 1 | 4.53 | 17.74 | 145.883 | 1 |
| Kynurenine | Kynurenine | C10H12N2O3 | [M+H] | 1 | 3.81 | 18.8 | 145.883 | 1 |
| Kynurenine | Kynurenine | C10H12N2O3 | [M+H] | 1 | 3.81 | 9.3 | 191.967 | 1 |
| Kynurenic acid | Kynurenic acid | C10H7NO3 | [M+H] | 1 | 4.83 | 19.41 | 143.883 | 1 |
| Kynurenic acid | Kynurenic acid | C10H7NO3 | [M+H] | 1 | 4.83 | 12.99 | 171.883 | 1 |
| Indole-3-acetic acid | Indole-3-acetic acid | C10H9NO2 | [M+H] | 1 | 5.98 | 15.36 | 129.967 | 1 |
| Indole-3-acetic acid | Indole-3-acetic acid | C10H9NO2 | [M+H] | 1 | 6 | 32.35 | 102.967 | 1 |
| DL-Indole-3-lactic acid | DL-Indole-3-lactic acid | C11H11NO3 | [M+H] | 1 | 5.67 | 11.93 | 159.967 | 1 |
| DL-Indole-3-lactic acid | DL-Indole-3-lactic acid | C11H11NO3 | [M+H] | 1 | 5.67 | 11.07 | 187.967 | 1 |
| 3-Indoleethanol | 3-Indoleethanol | C10H11NO | [M+H] | 1 | 6.06 | 13.59 | 143.883 | 1 |
| 3-Indoleethanol | 3-Indoleethanol | C10H11NO | [M+H] | 1 | 6.06 | 23.65 | 116.883 | 1 |

### Ex vivo system

The ex vivo system was modified from the Air-liquid interface^12^. In short, 10-14 week old mice (C57BL/6J) were kept on antibiotic water as previously described^3^ and then 50,000 pancreatic KPC cells (7940B) were injected into the pancreas. The orthotopic syngeneic pancreatic tumors were isolated from the mice 21 days post transplantation. Upon extraction, tumors were added to TIFM on ice and sliced within 5 min to about 2-3mm thickness using sterile scalpels before embedding in the ex vivo system. The bottom layer was created by combining 8 parts collagen (FujiFilm, Wako Chemicals, Cellmatrix Type I-A), with 1 part reconstitution buffer (2.2g NaHCO_3_ in 100 ml NaOH solution supplemented with 200mM HEPES) and 1 part 10x TIFM supplemented with 10% dialyzed FBS. The top layer was only collagen (FujiFilm, Wako Chemicals, Cellmatrix Type I-A). For the outside media, feeding the ex vivo system we used 1:1 TIFM to bacterial supernatant. Bacterial supernatant was generated by culturing *Klebsiella oxytoca*, *Klebsiella penumoniae* and *Klebsiella variicola* isolated in this study, as well as *E. coli K12*, as previously described in the metabolomics section. Since glucose was depleted in the supernatants collected from the bacteria, we supplemented the bacterial supernatant with 700 mg/l glucose (equivalent to glucose concentration in TIFM). The ex vivo system did not contain any antibiotics or antifungals, and the media was changed every 18 hours. Endpoints were at 22 and 44 hours, where the tissue was collected by fixing in Formalin-zinc solution for 24 hours before switching to 70% ethanol. The tissue was paraffin embedded and processed for histological evaluation. Fresh tumor tissues were collected during the embedding as a control for endpoint analysis.

### Immunofluorescence

Paraffin-embedded sections were rehydrated by washing slides twice with xylene, twice with 100% ethanol, and twice with 95% ethanol. Sections were then washed once with water to remove residues from previous washes. Slides were subsequently microwaved in CITRA Plus (BioGenex) for antigen retrieval. For preparation of use of the Alexa Fluor™ 488 Tyramide SuperBoost™ Kit (ThermoFisher), sections were incubated in 3% hydrogen peroxide for 15 min at room temperature (RT) to neutralize endogenous peroxidase activity. Sections were then blocked in 10% goat serum for 30 min at RT followed by incubation in primary antibody (Granzyme B [D2H2F], Cell Signaling) overnight at 4°C. The next day, sections were incubated in Poly-HRP-conjugated secondary antibody for an hour at RT followed by the Tyramide SuperBoost™ Amplification reaction as described in the kit’s protocol. Slides were once again microwaved in CITRA Plus and blocked in 1% BSA for 30 min at RT. Additional primary antibodies (CD8α [D4W2Z], Cell Signaling; E-Cadherin [4A2], Cell Signaling) were then added to the sections and incubated at 4°C overnight. The next day, sections were incubated in secondary antibody for 45 min at RT, counterstained for DAPI (1:30,000) for 7 min at RT, and mounted with Prolong Diamond Antifade Mountant (Invitrogen). Images were taken using Olympus BX53F microscope of at least 3 random non-overlapping fields and quantified using ImageJ to measure percentage of positive area and percentage of overlapping positive areas.

### Immunohistochemistry

Murine pancreatic tissue grown ex vivo was as described above (see ex vivo system). Sections of the tissue were used for immunohistochemistry using anti-Ki67 antibody (Abcam, ab15580 Lot:1063776-1) diluted 1:500. Stained slides were analyzed using positive cell detection in Qupath v0.4.0^13^.

### Statistical analysis

Statistical analyses were performed using PRISM 10 for macOS, Version 10.4.0 (527), October 23, 2024.

## Results

### Isolation of bacteria and fungi from human pancreatic tumors and small intestine

Human pancreatic small intestine and tumor tissue samples were collected from patients at the University of Michigan undergoing surgical resection with prior consent (**Table 1**). For the isolation of live microbiota, we were able to use cryopreserved tissue that was either directly flash frozen or frozen in sterile RPMI media, for up to two years after collection, when stored at -80°C **(Figure 1).** We chose a variety of media composition to enrich for fungi and bacteria (**Table 2**). These included routinely used media conditions modified to adjust, for example, for the higher pH found in the pancreas or supplemented with nutrients to enrich for recently published long-term survivor associated bacteria^6^. After the initial isolation step, we then isolated pure colonies from the mixed colony plates grown under hypoxic or anoxic conditions. While we aimed to use the same amount of tissue for the isolation per plate, colony forming units varied greatly between patients. Notably, this was not a predictor for microbial diversity as seen when comparing patients A and B (**Table 3 and 4**). We were able to isolate a variety of gram negative and gram positive bacteria, but were not able to isolate any of the long term survivor associated bacteria such as *Pseudoxanthomonas, Saccharopolyspora* or *Streptomyces*^6^ (**Table 4**). Since fungi are often overlooked in microbiome studies, we also included media conditions to enrich for fungi, such as the Sabaroud agar, and successfully isolated several fungi such as *Candida* and *Saccharomyces*. In the early optimization process, we also tested FastFung^14^ media but we were not able to isolate fungi in hypoxic or anoxic conditions. When we applied our pipeline to healthy donor pancreatic tissue (from organ/tissue donors through Gift of Life Michigan), we also included the Leeming-Notman agar and were able to isolate *Candida* and *Malassezia*. Since *Malassezia sp.* is not currently included in the database of the MALDI-TOF used for this study, we grew all unidentified colonies from the Leeming-Notman onto the CHROMagar™ Malassezia and identified one isolate as *Malassezia globosa* (**Supplemental Figure 1**). Of note, when tissue is streaked directly onto the CHROMagar™ Malassezia we did not observe any fungal growth, therefore the application of Leeming-Notman agar provides an important substrate for initial fungal growth from tissue isolation.

**Figure 1:**
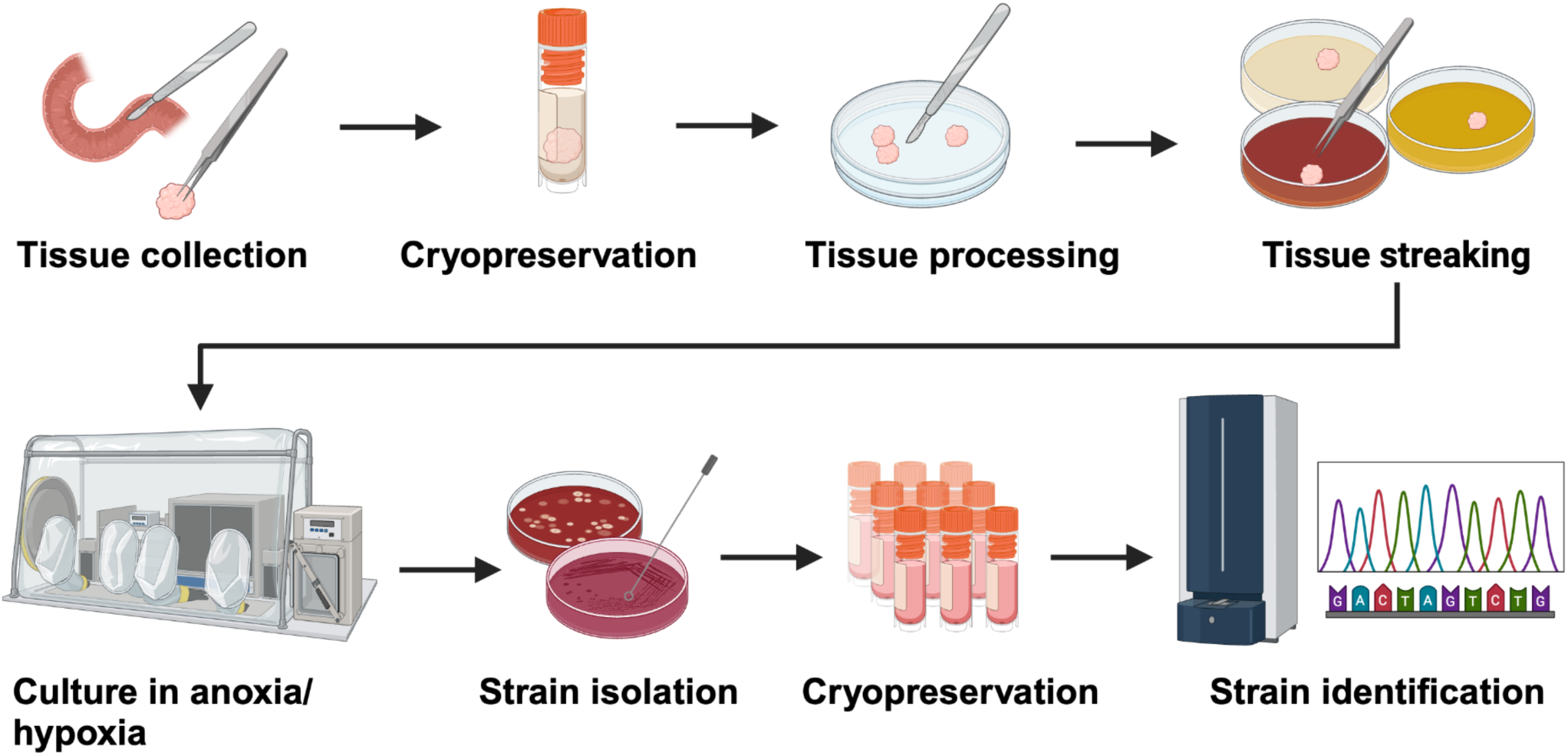
Scheme of the isolation process of bacteria and fungi from the small intestine and tumor of pancreatic cancer patients.

**Table 1:** Tissue used for the isolation.

| Patient | Sample | Pathology |
| --- | --- | --- |
| A | Small intestine | Pancreatic cancer patient |
| B | Small intestine | Pancreatic cancer patient |
| C | Small intestine | Pancreatic cancer patient |
| D | Small intestine | Pancreatic cancer patient |
| E | Small intestine and tumor | Pancreatic cancer patient |
| F | Small intestine and pancreas | Organ donor |

**Table 2:** Media composition used for the study.

| Name | Media | Modification | Selection for |
| --- | --- | --- | --- |
| M1 | Sabourad Agar | No antibiotics | Fungi and bacteria |
| M2 | BHI Agar | CaCl<sub>2</sub> | Bacteria, i.p. *Streptomyces* |
| M3 | Bacillus clausii Agar | pH 8.5 | Bacteria, i.p. *B. clausii* and simulation of pH in pancreas |
| M4 | Columbia Agar | NaCl, tryptophan, histidine, NaNO<sub>2</sub> | Bacteria, i.p. *Pseudoxanthomonas*, *Saccharopolyspora* |
| M5 | GAM Agar |  | Anerobic bacteria |
| M6 | LN Agar | No antibiotics | Fungi, in particular lipid dependent |

**Table 3:** Colony forming units (CFU) per plate. The colony forming units were monitored daily for up to 21 days. Legend: **Small int:** small intestine **n.i:** media not included in this isolation; **\*** bacterial overgrowth within 2-3 days.

| Patient | Tissue | Hypoxia M1 | Hypoxia M2 | Hypoxia M3 | Hypoxia M4 | Hypoxia M5 | Hypoxia M6 |
| --- | --- | --- | --- | --- | --- | --- | --- |
| A | Small int. | 10 | 71 | 21 | 40 | 0 | n.i. |
| B | Small int. | 29 | 59 | 5 | 58 | 21 | n.i. |
| C | Small int. | 5 | 0 | 0 | 2 | 3 | n.i. |
| D | Small int. | * | 6 | 5 | * | 14 | n.i. |
| E | Small int. | 12 | 1 | 2 | 18 | 7 | n.i. |
| F | Small int. | 45 | 105 | 69 | 54 | 70 | 123 |
| Patient | Tissue | Anoxia M1 | Anoxia M2 | Anoxia M3 | Anoxia M4 | Anoxia M5 | Anoxia M6 |
| A | Small int. | 9 | 9 | 72 | 180 | 12 | n.i. |
| B | Small int. | 35 | 4 | 16 | 11 | 5 | n.i. |
| C | Small int. | 0 | 0 | 0 | 1 | 2 | n.i. |
| D | Small int. | 26 | 6 | 21 | 12 | 9 | n.i. |
| E | Small int. | 5 | 0 | 2 | 1 | 0 | n.i. |
| F | Small int. | 25 | 27 | 11 | 19 | 21 | 8 |
| Patient | Tissue | Hypoxia M1 | Hypoxia M2 | Hypoxia M3 | Hypoxia M4 | Hypoxia M5 | Hypoxia M6 |
| E | Tumor | 32 | 16 | 13 | 18 | 4 | n.i. |
| F | Pancreas | 19 | 13 | 35 | 29 | 14 | 0 |
| Patient | Tissue | Anoxia M1 | Anoxia M2 | Anoxia M3 | Anoxia M4 | Anoxia M5 | Anoxia M6 |
| E | Tumor | 9 | 12 | 2 | 4 | 0 | n.i. |
| F | Pancreas | 1 | 4 | 0 | 2 | 3 | 0 |

**Table 4:**
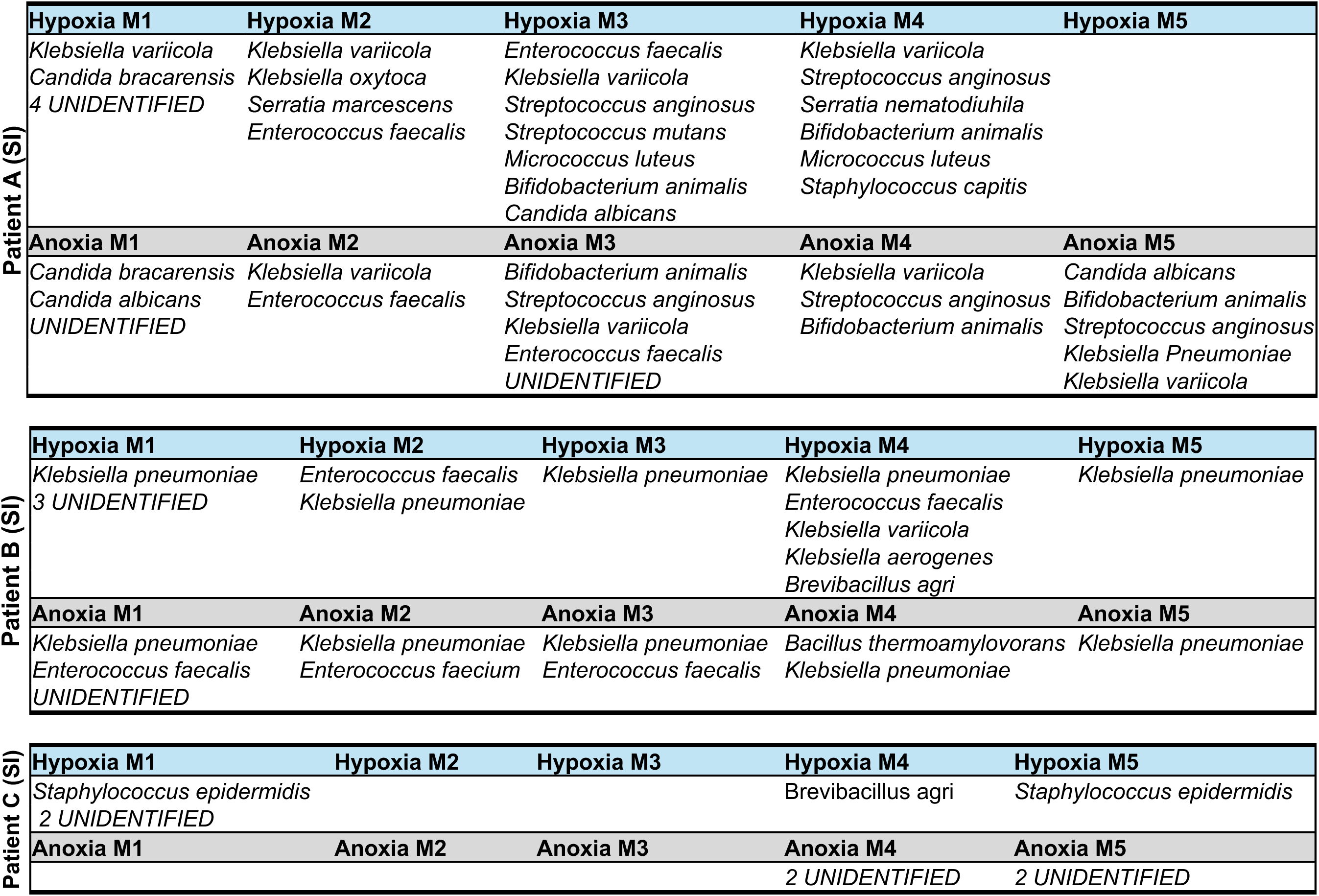
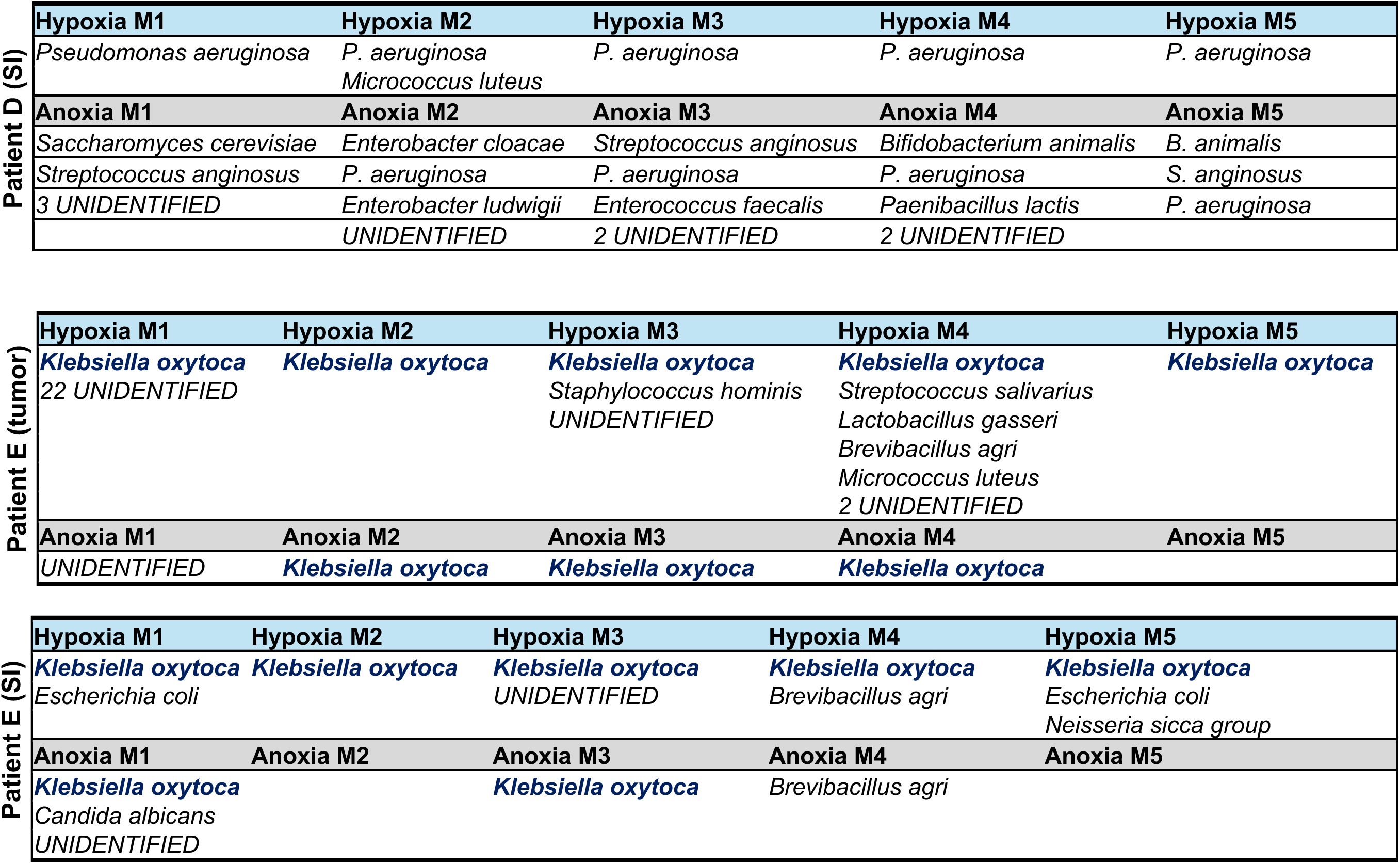
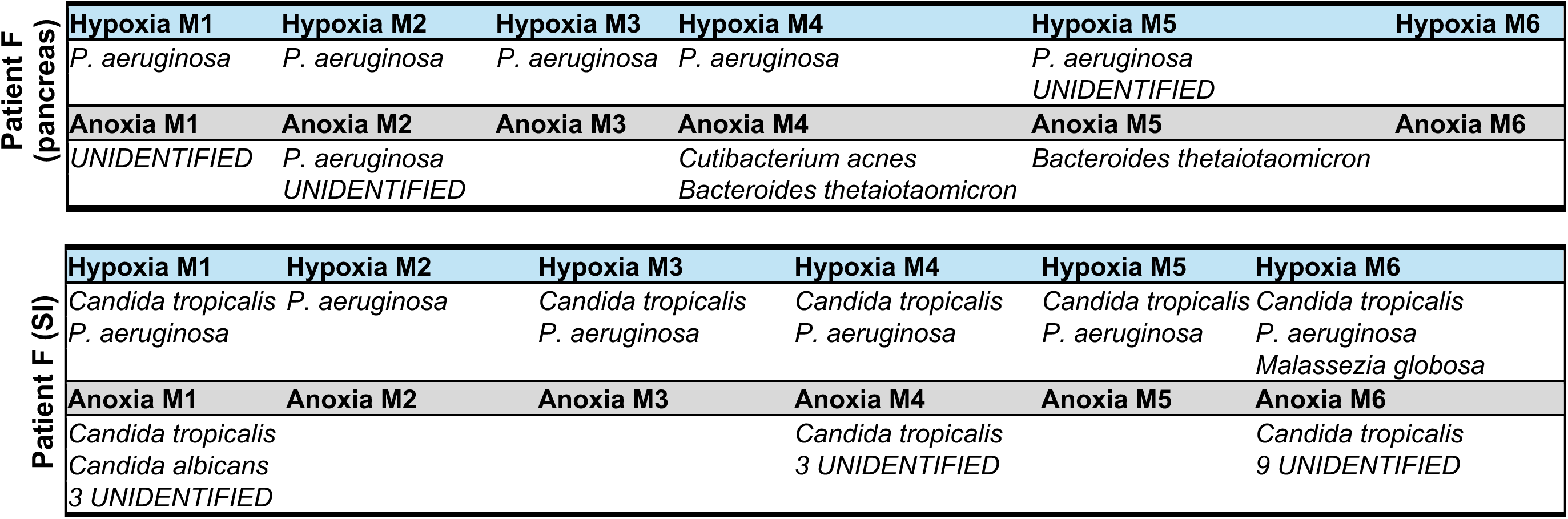
Isolated bacteria and fungi from small intestine (SI) and tumor.

All isolations were performed in hypoxic and anoxic conditions to simulate the small intestine and pancreatic tumor environment, which in turn also prevented overgrowth by bacteria. We found that if plates were grown in normoxia, fungal isolation was not possible due to the overgrowth of bacteria. Based on our MALDI-TOF analysis, we did not isolate fungi from the human pancreatic tumor but isolated several bacteria such as *Streptococcus salivarius, Lactobacillus gasseri, Brevibacillus agri, and Micrococcus luteus.* However, the most abundant bacteria found in the pancreatic tumor based on our isolation was *Klebsiella oxytoca*. This trend was also seen in previous studies that applied a sequencing approach ^2,3^. Based on the literature and the dominance of *Klebsiella oxytoca* in our isolated cultures, we decided to focus on this strain for further characterization.

### Genomic sequencing of Klebsiella oxytoca reveals several genes that are associated with treatment resistance

To learn more about *K. oxytoca* strain that we isolated from the human pancreatic tumor, herein designated as *UMKO1,* we performed bacterial genome sequencing (**Figure 2A, Supplemental table 1**). The sequencing results identified a contig of 6,070,641 bp, confirming the identity of the strain as *Klebsiella oxytoca* (NZ_KQ088007.1 Klebsiella oxytoca strain CHS143 aetza-supercont1.1, **Figure 2B**). The second contig was identified as the plasmid pCAV1335-115 (Klebsiella oxytoca strain CAV1335 plasmid pCAV1335-115, **Figure 2B**). We then compared our sequencing data to several publicly available *Klebsiella oxytoca* sequencing data sets (**Supplemental table 2**) using the Pan-genome explorer^11^. We found several gene clusters that are unique to the newly identified *Klebsiella oxytoca* strain (**Figure 2C**), which included a wealth of metabolic genes **(Supplemental table 3)**. In addition, our analysis suggests that the isolated *K. oxytoca* strain *UMKO1* contains a predicted protein sequence for cytidine deaminase. Given the role of the long version of cytidine deaminase in gemcitabine resistance in pancreatic cancer^2^, we compared the predicted amino acid sequence from our clinical isolate to the recently published amino acid sequence^2^ and confirmed that the isolated *K. oxytoca UMKO1* strain indeed possesses the long version of cytidine deaminase (**Sup figure 2**).

**Figure 2:**
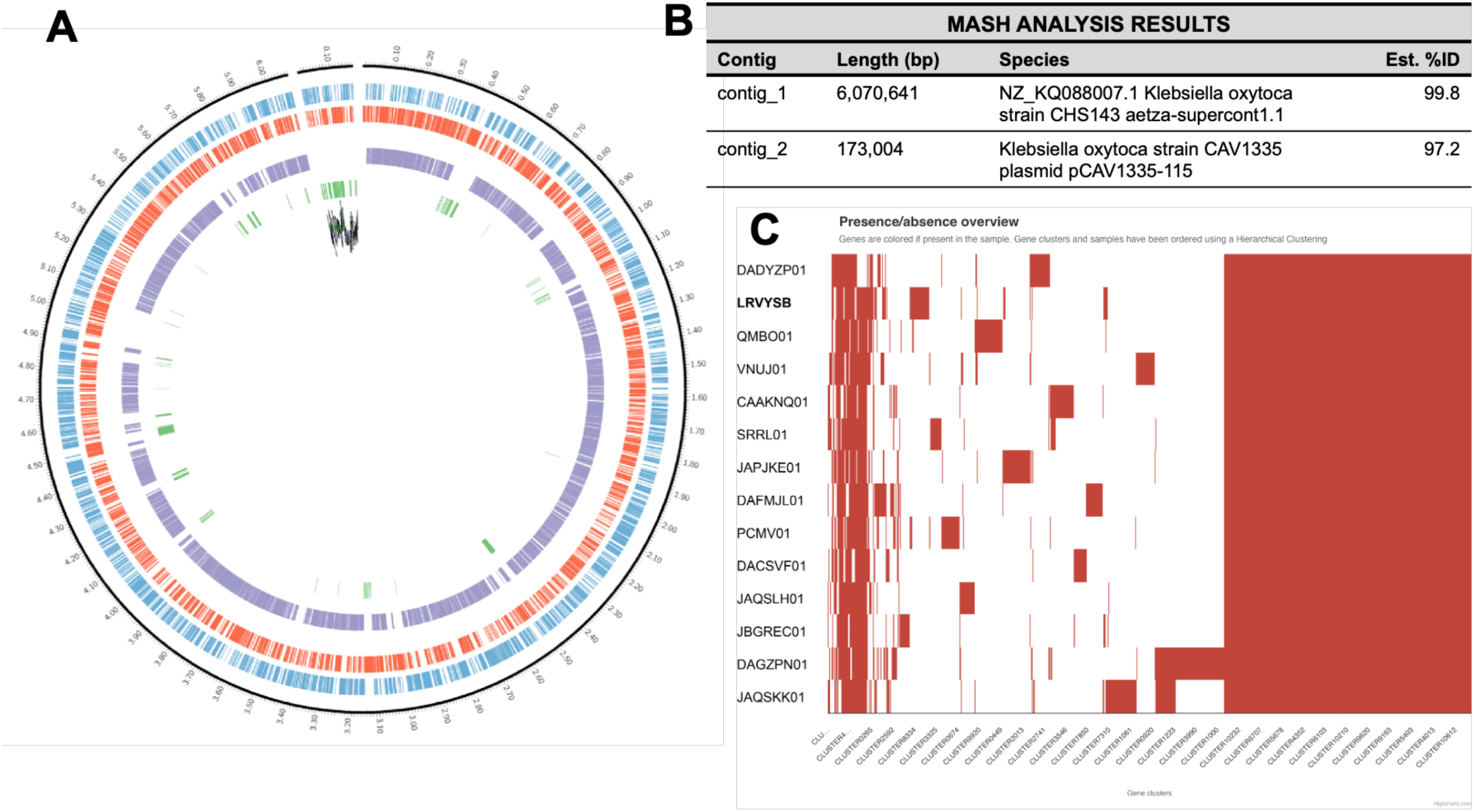
Sequence analysis of *Klebsiella oxytoca (UMKO1),* isolated from a human pancreatic tumor. **(A)** Circos plot of the bacterial genomic sequence of the isolated *Klebsiella oxytoca UMKO1* using the Pan-genome explorer tool. Forward genes are in blue, reverse genes in red, core genes in purple and strain specific genes in green. Contig 2 displays a GC skew. **(B)** MASH analysis results show two contigs, a match to *Klebsiella oxytoca* strain CHS143 and to the plasmid pCAV1335-115. **(C)** Gene presence/absence analysis shows a distinct cluster of genes unique to the isolated *Klebsiella oxytoca* strain *UMKO1*, the full list can be found in supplemental table 3.

### Klebsiella oxytoca produces indoles in the tumor microenvironment that are associated with the reprogramming of the tumor microenvironment and therapy outcome

Recent studies demonstrated that microbial-derived metabolites contribute to the immunosuppressive pancreatic tumor microenvironment and can interfere with the patient’s response to therapy^15–17^. To determine which microbial metabolites are derived by *Klebsiella oxytoca* in the tumor microenvironment, we grew the strain *UMKO1* in Tumor Interstitial Fluid Media (TIFM)^18^ and performed untargeted metabolomics of polar metabolites using LC-MS/MS (**Figure 3**). The use of TIFM allowed us to determine the production of bacterial derived metabolites by *UMKO1* in a physiological relevant nutrient environment, simulating the nutrients available in the pancreatic TME^18^. We found that *Klebsiella oxytoca* depleted several amino acids from the media, including tryptophan (**Figure 3, Sup figure 3A**). We also observed other key metabolites, such as alpha ketoglutarate and citric acid, known to be utilized by bacteria, were depleted from the media (**Sup Figure 3B).** In addition, our untargeted analysis suggested the production of polyamines such as putrescine **(Sup Figure 3C)** and the production of butyrate **(Sup Figure 3D).** Given the depletion of tryptophan and the production of various indoles, we then asked if the breakdown of tryptophan could be leading to the production of clinical relevant indoles, such as indole-3-acetic acid, when grown in TIFM. We first observed production of indole-3-acetic acid in the untargeted analysis (**Figure 3C**). In support, the bacterial genomic sequence analysis revealed several enzymes associated with the indole metabolic pathway, such as *tryptophanase* and *indole pyruvate decarboxylase*. To confirm our untargeted analysis we applied a newly developed, custom targeted LC-MS/MS method to quantify tryptophan and selected metabolites in the kynurenine and indole pathway. Our results demonstrate that *UMKO1* can produce not only key indoles such as indole-3-acetic acid and 3-indole ethanol, but also kynurenic acid (**Figure 3**). Kynurenic acid and indole-3-acetic acid have been identified in previous studies as potent aryl hydrocarbon receptor agonists, which can reprogram macrophages into immunosuppressive tumor associated macrophages^16^.

**Figure 3:**
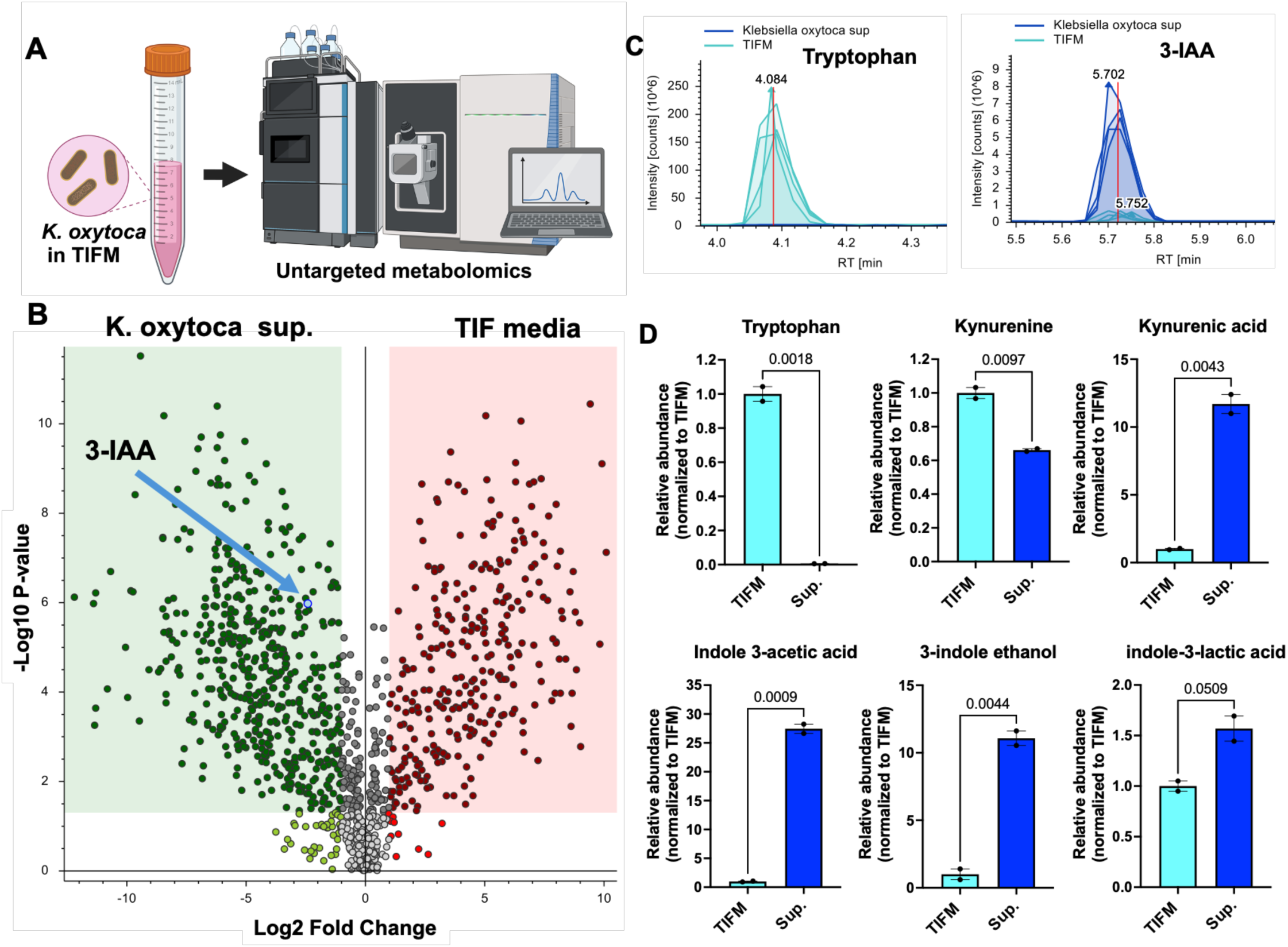
*Klebsiella oxytoca UMKO1* produces indole-3-acetic acid and other tryptophan derived metabolites when grown in tumor nutrient like environment. **(A)** *Klebsiella oxytoca UMKO1* was grown in tumor interstitial fluid media (TFIM) and polar metabolites were extracted for untargeted metabolomics using LC-MS/MS. **(B)** Volcano plot of the identified metabolic features enriched in the bacterial supernatant or TIFM (3-IAA: indole 3-acetic acid). **(C)** Tryptophan was depleted in the bacterial supernatant and **(D)** tryptophan derived metabolites were enriched. Statistical analysis: unpaired t-test, P value as indicated on graph.

### Klebsiella oxytoca reprograms the pancreatic tumor microenvironment

To test how the pancreatic tumor microenvironment is influenced by bacterial metabolites produced by intratumoral bacteria, we grew several of the isolated bacterial strains in TIFM overnight and collected their supernatant. These conditioned media were applied to murine pancreatic tumors grown ex vivo in physiological relevant nutrients (TIFM) containing bacterial supernatants for up to 44 hours. We found no differences in CD8 T cell abundance in the tumors exposed to the TIFM control or bacterial supernatants (**Figure 4A,B**). However, we observed a decrease in Granzyme B-positive CD8 T cells in the tumor explants when grown in the presence of *UMKO1* supernatant, suggesting decreased cytotoxic T cell activity. We saw the opposite trends when tumors were treated with the supernatant from *Klebsiella pneumonia* or the standard *E. coli K12*, each grown in TIFM (**Figure 4**). In addition, we found that the addition of *K. oxytoca* strain *UMKO1* supernatant increased the overall stain for Ki67 in the ex vivo system, suggesting an increase in proliferation of cells in the tumor microenvironment in the presence of *UMKO1* (**Sup Figure 4**). Together, these data illustrate potential ways in which the supernatant of *K. oxytoca* can uniquely contribute to an immunosuppressive environment in pancreatic cancer.

**Figure 4:**
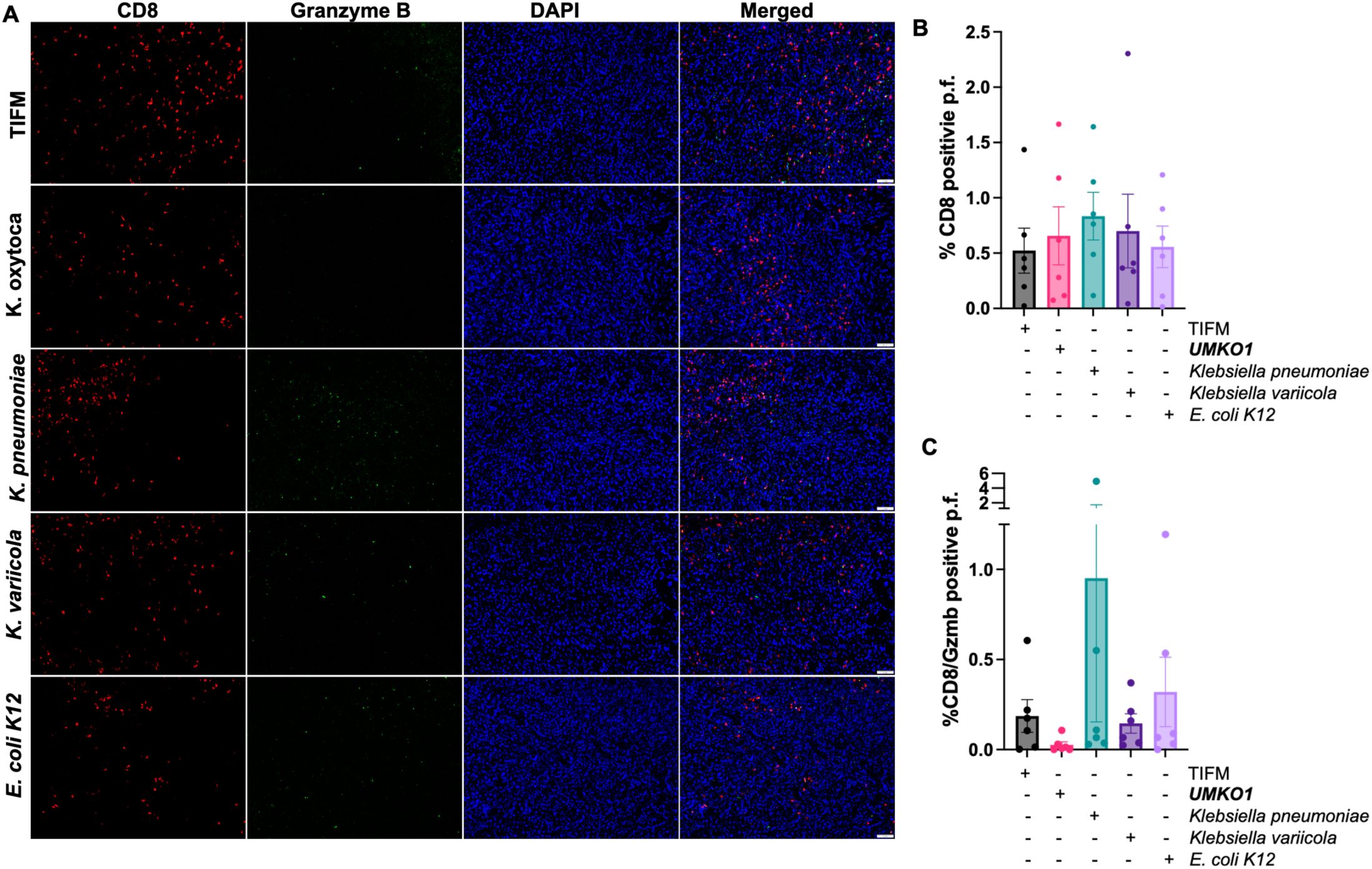
*Klebsiella oxytoca UMKO1* supernatant reduces granzyme B positive CD8 T cells in an ex vivo tumor system. **(A)** Representative images of tumor explants grown ex vivo for 44 hours, supplemented with TIFM or various bacterial condition media (TIFM + bacterial supernatant, 1:1). **(B)** CD8 T cell abundance does not change with bacterial supernatant but **(C)** granzyme B positive T cells were reduced in tumor explants exposed to *Klebsiella oxytoca UMKO1* supernatant. Two murine pancreatic tumor explants were quantified for CD8 and granzyme B staining, and granzyme B positive CD8 T cells were reported in **(C)**. Three images per condition, graphed in **(B)** and **(C)**. The staining and the quantification were performed blinded to avoid any potential bias.

**Figure 5:**
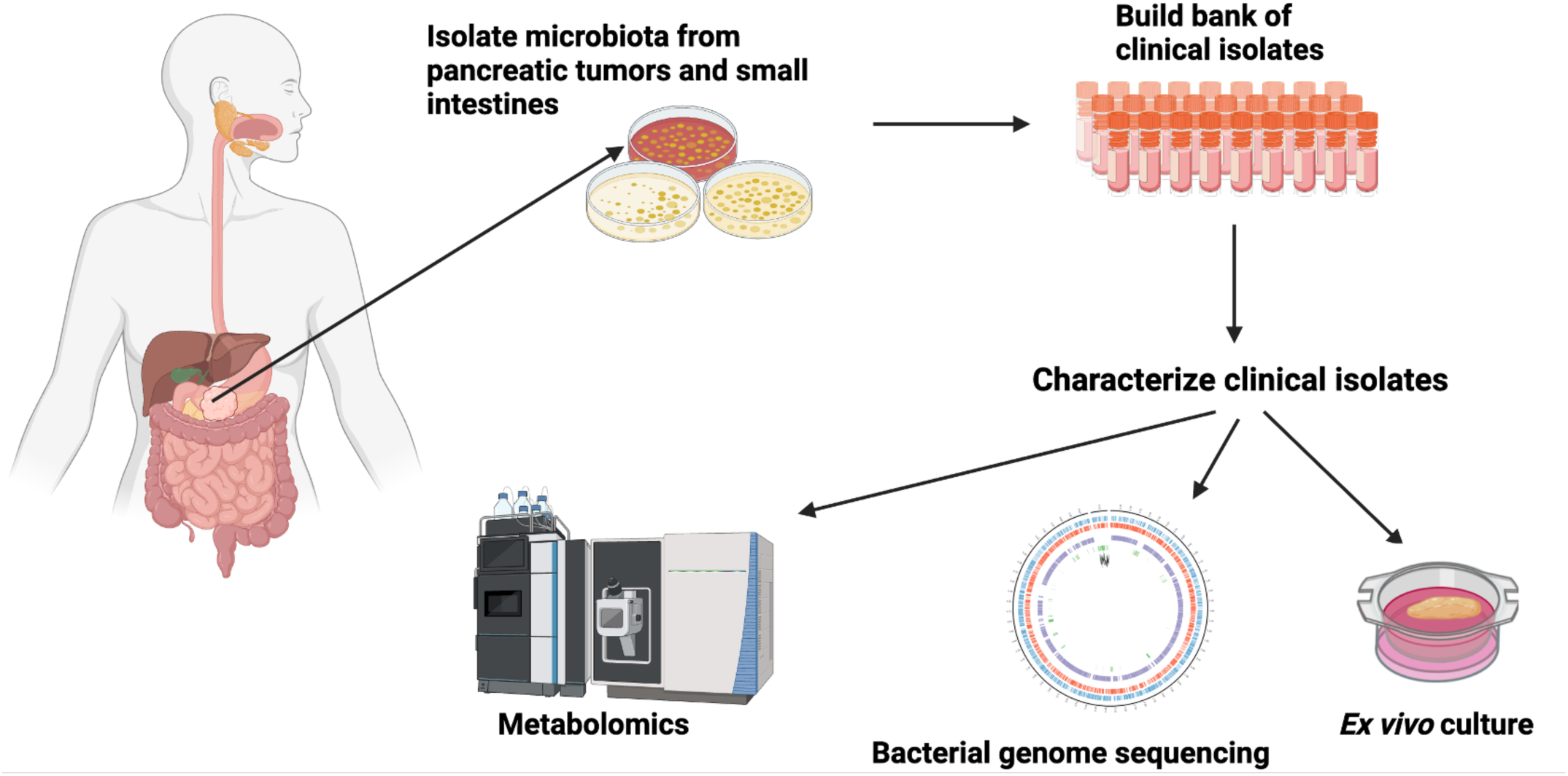
Graphical abstract.

## Discussion

One of the major obstacles when studying the pancreatic tumor microbiome is its low biomass. While bacteria and fungi can be identified in the tumor by immunohistochemistry ^2–5^, the low abundance can make it difficult to interpret sequencing analysis due to the high noise to signal ratio. Current functional studies are often limited to FMTs, or bacteria and fungi that are isolated from sources other than pancreatic cancer patients. To address this, we started a collection of bacteria and fungi isolated from pancreatic cancer patients at the University of Michigan. We found that we were able to isolate live microbiota from flash frozen tissue, with or without RPMI, up to two years after the initial collection. Our media composition was optimized to enrich for fungi, especially yeast, which is often found in the small intestine and has been reported in the pancreatic tumor microenvironment^3,5^. In addition, we also tried to enrich for bacteria found in previous sequencing data, including long term survivor associated bacteria^6^. While we were not able to isolate any long-term survivor associated bacteria, we successfully isolated a wealth of bacteria from the small intestine and tumor of pancreatic cancer patients. In contrast we only isolated fewer unique strains from the donor pancreas, and only one Gammaproteobacteria, *Pseudomonas aeruginosa*. Interestingly, while we found a total of 110 colony forming units (CFU) across all hypoxic plates from the donor pancreas isolation, all but one of these colonies were identified as *P. aeruginosa.* Given the rapid growth of *P. aeruginosa* observed on the plates, which had bacterial overgrowth in a matter of days under hypoxic conditions, we conclude the high amount of CFUs is due to the rapid growth of *P. aeruginosa* and not a reflection of higher microbial abundance in the donor pancreas. This overgrowth by *P. aeruginosa was* also seen in other tissues used for this study (**Table 3, 4;** Patient D, M1, M4). In addition, we isolated *Cutibacterium acnes* and *Bacteroides thetaiotaomicron* from the human donor pancreatic tissue when grown in anoxic conditions, which delayed the bacterial overgrowth by *P. aeruginosa*. Importantly, we did not see any bacterial or fungal growth when we plated the PBS solution that was used to rinse the instruments used for dissection of the pancreas and small intestine tissue. This suggests that the microbes isolated from our pancreas and small intestine tissues are unlikely to be contaminants introduced during tissue dissection.

The most abundant bacteria found in the pancreatic tumor was *Klebsiella oxytoca*. We confirmed the presence of the long form of the cytidine deaminase and unique metabolic enzymes involved in tryptophan metabolism in the bacterial genomic sequence of our *UMKO1* strain*. UMKO1* is also able to produce indole-3-acetic acid when grown in TIFM. Furthermore, we observed *UMKO1’s* ability to reprogram the murine pancreatic tumor microenvironment, as evidenced by the decrease of granzyme B positive CD8 T cells in the presence of *Klebsiella oxytoca* supernatant. These data support the potential pro-tumorigenic role of *K. oxytoca UMKO1* in the PDAC tumor microenvironment and corroborate findings previously reported previously ^2,3^.

The microbiota isolated and characterized in this study, will aid in providing useful resources for mechanistic studies to gain novel insights into the complex role of the microbiome in pancreatic cancer.

## Author contribution

D.A., H.A. and D.D. conceptualized this study; D.A. prepared the first draft of this manuscript, which was then edited by H.A., R.H., T.M.S, D.D. and C.A.; D.A., H.A., R.H. K.K. lead the experimental work of the isolation under the guidance of T.M.S and D.D.; M.H., and J.A.N.C. helped with the initial optimization of the isolation workflow and early MALDI-TOF analysis; C.S. performed the tissue processing for histology; M.PM. and T.F. guided the collection of the pancreata donor sample; D.A, and L.Z. performed the metabolomics analysis under the guidance of C.A.L.; D.A., N.N. and R.H. performed the ex vivo system studies and immunohistochemistry analysis; A.B. performed the IF staining of the ex vivo system; A.dD. and D.A. performed the analysis of the bacterial genome sequencing results.

## Financial support

This work was supported with financial support from the NIH (K08CA282374), the Robert Wood Johnson Foundation, and the American College of Surgeons (D.D). D.A. received additional support from the University of Michigan Postdoctoral Pioneer Program and the NIH (NIAID Training Grant T32-AI007413). H.A. received support from Rogel Cancer Center, NIH (CBTP-T32 CA009676 and T32TR004371). A.B. received financial support from the NIH (T32-DK094775 and 1F31CA284505-01A1). C.A.L. was supported by the NCI (R37CA237421, R01CA248160, R01CA244931) and NHLBI (P01 HL149663).

## Declaration of interests

C.A.L. has received consulting fees from Odyssey Therapeutics and is an inventor on patents pertaining to Kras-regulated metabolic pathways, redox control pathways in pancreatic cancer, and targeting the GOT1-pathway as a therapeutic approach (US Patent No: 2015126580-A1, 05/07/2015; US Patent No: 20190136238, 05/09/2019; International Patent No: WO2013177426-A2, 04/23/2015). D.D. is a co-inventor on patents pertaining to immune cell targets for therapy in solid tumors (US Patent No: 2018044422-A1, 07/31/2017; US Patent No: 2018023111-A1, 07/31/2017).

## Acknowledgment

We want to thank first and foremost the patients that made this study possible. We would like to thank the University of Michigan Microbiome core facility for the MALDI-TOF analysis. We also would like to thank Alex Muir for supplying the Tumor Interstitial Fluid Media (TIFM) and helpful discussion regarding the use of the media. We would also like to thank the members of the Rogel and Blondy Pancreatic Cancer Center at the University of Michigan, as well as all the members of the Daley, Lyssiotis and Schmidt lab for their helpful suggestions and feedback. Figures were created in part with Biorender.

