## Supplemental figures for "Isolation and characterization of microbiota from human pancreatic tumors and small intestine"

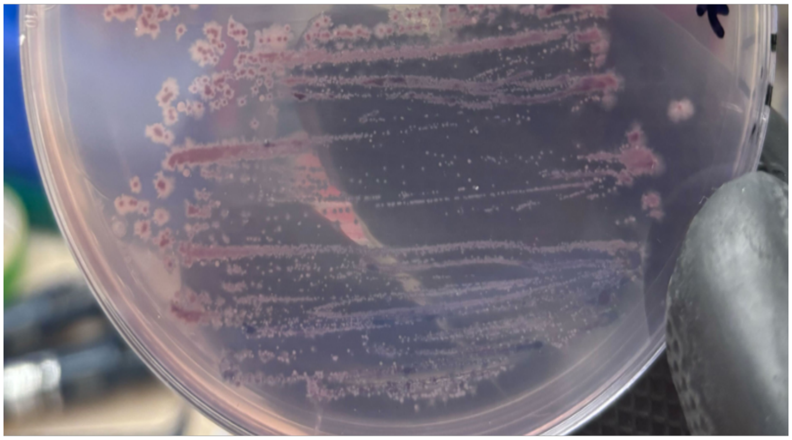

**Supplemental Figure 1: Unidentified fungus from Patient F grown on ChromAgar Malassezia**

**Supplemental Table 1: Additional information on the sequencing results shown in Figure 2**

| Statistics | |
|---|---|
| Total bp sequenced | 722,506,331 bp |
| Total number of reads | 204,426 reads |
| Longest read | 76,682 bp |
| Raw coverage | 115x |
| Assembled coverage | 104x |
| Genome size (Mb) | 6.2 Mb |
| Number of contigs | 2 contigs |
| Number of genes annotated | 5,944 genes |

**Supplemental Table 2: *Klebsiella oxytoca* strains used for the comparison in Figure 2C.**

| Accession Number | Strain/Isolate | Location | Region | Source species | Source material |
|---|---|---|---|---|---|
| **LRVYSB** | ***UMKO1*** | **USA** | **Michigan** | **Homo sapiens** | **Pancreatic tumor** |
| DAGZPN01 | F58220114 | USA | Utah | NA | Wastewater |
| DAFMJL01 | | Spain | Unknown | Homo sapiens | |
| DACSVF01 | ARLG-4836 | USA | South | Homo sapiens | Urine |
| DADYZP01 | 1D-1188 | Denmark | Unknown | Homo sapiens | |
| CAAKNQ01 | MC1 | USA | Unknown | Homo sapiens | Feces |
| JAQSLH01 | HD2641 | China | Chengdu | Homo sapiens | Respiratory tract |
| JAQSKK01 | HD8859 | China | Suzhou | Homo sapiens | Abdominal cavity |
| JBGREC01 | Kox-70 | India | Assam | Labeo rohita (Rohu Carp) | |
| VNUJ01 | 708994-2009 | Switzerland | Unknown | Homo sapiens | Urine |
| PCMV01 | UNM | USA | New Mexico | Homo sapiens | blood |
| JAPJKE01 | KM57/09 | Switzerland | Unknown | Equus caballus | Pus |
| SRRL01 | BX1S15 | USA | Philadelphia | NA | Retail fresh cilantro |
| QMBO01 | JK01 | USA | Rhode Island | Mus musculus | Spleen |

**Supplemental Table 3: Unique gene clusters identified in *UMKO1,* when compared to other *Klebsiella oxytoca* strains**

| Cluster | Gene | Cluster | Gene |
|---|---|---|---|
| 9802 | Tn3 family transposase TnEc1 | 2250 | Aliphatic nitrilase |
| 9801 | DNA-invertase hin | 2247 | 4-hydroxy-tetrahydrodipicolinate synthase |
| 7980 | Sec-independent protein translocase protein TatA | 2066 | L-threonine ammonia-lyase |
| 7749 | Riboflavin transporter RfnT | 2064 | Glutathione-binding protein GsiB |
| 7745 | ATP-dependent DNA helicase Rep | 2063 | 8-oxoguanine deaminase |
| 7355 | HTH-type transcriptional regulator CdhR | 2061 | Protein UmuC |
| 6719 | IS110 family transposase ISSm1 | 2058 | 4-hydroxy-tetrahydrodipicolinate synthase |
| 6718 | RepFIB replication protein A | 1959 | IS66 family transposase ISKpn24 |
| 5847 | Cyclic UMP-AMP synthase | 1952 | Porin OmpL |
| 5837 | GTP 38-cyclase | 1746 | IS1 family transposase IS1R |
| 5497 | Cytochrome bo3 ubiquinol oxidase subunit 3 | 1542 | IS3 family transposase ISKpn38 |
| 5088 | cGAMP-activated phospholipase | 1541 | Acetyl-coenzyme A synthetase |
| 5087 | Cytochrome bo3 ubiquinol oxidase subunit 4 | 1328 | IS3 family transposase ISKpn38 |
| 4540 | ISNCY family transposase ISLad2 | 1327 | HTH-type transcriptional regulator ArgP |
| 4539 | S-formylglutathione hydrolase FrmB | 1326 | Carnitine transport ATP-binding protein OpuCA |
| 4536 | Cyclic GMP-AMP synthase | 1316 | Tyrosine recombinase XerC |
| 4525 | RNA polymerase-binding transcription factor DksA | 1313 | 45-DOPA dioxygenase extradiol |
| 3865 | Glutamine transport ATP-binding protein GlnQ | 1105 | Protease HtpX |
| 3863 | Glutathione transport system permease protein GsiC | 1101 | IS3 family transposase ISEam1 |
| 3862 | Vitamin B6 salvage pathway transcriptional repressor PtsJ | 1097 | Putative dimethyl sulfoxide reductase chain YnfE |
| 3858 | DNA adenine methylase | 1093 | Tyrosine recombinase XerC |
| 3673 | Glutathione transport system permease protein GsiD | 1085 | 45-DOPA dioxygenase extradiol |
| 3480 | Beta-peptidyl aminopeptidase BapA | 867 | Outer membrane protein OprM |
| 3474 | DNA adenine methylase | 861 | 7-cyano-7-deazaguanine synthase |
| 3289 | D-aminopeptidase | 859 | Sodium-dependent dicarboxylate transporter SdcS |
| 3081 | HTH-type transcriptional regulator CysB | 657 | 3-[3aS4S7aS-7a-methyl-1 5-dioxo-octahydro-1H-inden-4-yl]propanoyl:CoA ligase |
| 3079 | Tyrosine recombinase XerD | 652 | putative multidrug resistance protein EmrK |
| 3068 | putative metal-dependent hydrolase YcfH | 650 | Membrane-bound lytic murein transglycosylase F |
| 2881 | Thiamine-monophosphate kinase | 642 | Sodium-dependent dicarboxylate transporter SdcS |
| 2668 | IS110 family transposase ISSaen1 | 641 | ATP-dependent DNA helicase Rep |
| 2460 | L-Ala-D/L-Glu epimerase | 410 | Multidrug export protein EmrB |
| 2459 | Protein Ves | 149 | IS3 family transposase ISYps8 |
| 2254 | Delta1-pyrroline-2-carboxylate reductase | 147 | cUMP-AMP-activated phospholipase |
| 2252 | Glutathione import ATP-binding protein GsiA | 146 | Cytochrome bo3 ubiquinol oxidase subunit 2 |
| 2251 | HTH-type transcriptional repressor NagR | 144 | Inner membrane protein YagU |

| Score | Expect | Method | Identities | Positives | Gaps |
| --- | --- | --- | --- | --- | --- |
| 583 bits(1502) | 0.0 | Compositional matrix adjust. | 284/294(97%) | 291/294(98%) | 0/294(0%) |

```
CDD_L       MHPRFQAVLPQLAADLQAAIAPMLADPHFPALLNADQVAALQSATGLDEDALAFALLPLA  60
            MHPRFQAVLPQLAADLQAAIAPMLADPHFPALLNADQVAALQSATGLDEDALAFALLPLA
K. oxytoca  MHPRFQAVLPQLAADLQAAIAPMLADPHFPALLNADQVAALQSATGLDEDALAFALLPLA  60

CDD_L       AACARTDLSHFNVGAIARGISGTWYFGGNMEFLGATMQQTVHAEQSAISHAWLRGEKSLL  120
            AACAR DLSHFNVGAIARGISGTWYFGGNMEFLGATMQQTVHAEQSAISHAWLRGEKSL
K. oxytoca  AACARADLSHFNVGAIARGISGTWYFGGNMEFLGATMQQTVHAEQSAISHAWLRGEKSLR  120

CDD_L       AITVNYTPCGHCRQFMNELNSGLALRIHLPGREAHSLQHYLPDAFGPKDLNIKTLLMDEQ  180
            AITVNYTPCGHCRQFMNELNSGL+LRIHLPGREAHSLQHYLPDAFGPKDL+IKTLLMDEQ
K. oxytoca  AITVNYTPCGHCRQFMNELNSGLSLRIHLPGREAHSLQHYLPDAFGPKDLDIKTLLMDEQ  180

CDD_L       NHGFPLSGDALAQAAIQAANRCHMPYSHSPSGVALELKDGTIFTGSYAENAAYNPTLPPL  240
            +HGFPLSGDALAQAAIQAANRCHMPYS+SPSGVALELKDGT+FTGSYAENAA+NPTLPPL
K. oxytoca  DHGFPLSGDALAQAAIQAANRCHMPYSNSPSGVALELKDGTLFTGSYAENAAFNPTLPPL  240

CDD_L       QGALNLLSLNGYDYPDIQRAVLAEKADAALIQWDATAATLKALGCHNIDRVLLG  294
            QGALNLLSLNGYDYPDIQRA+LAEKADAALIQWDAT ATLKALGCHNIDRVLLG
K. oxytoca  QGALNLLSLNGYDYPDIQRAILAEKADAALIQWDATVATLKALGCHNIDRVLLG  294
```

**Supplemental Figure 2: Predicted protein sequence of the cytidine deaminase identified in the newly isolated K. oxytoca strain compared to the long form of cytidine deaminase publis [illegible]y Geller et al, 2017.**

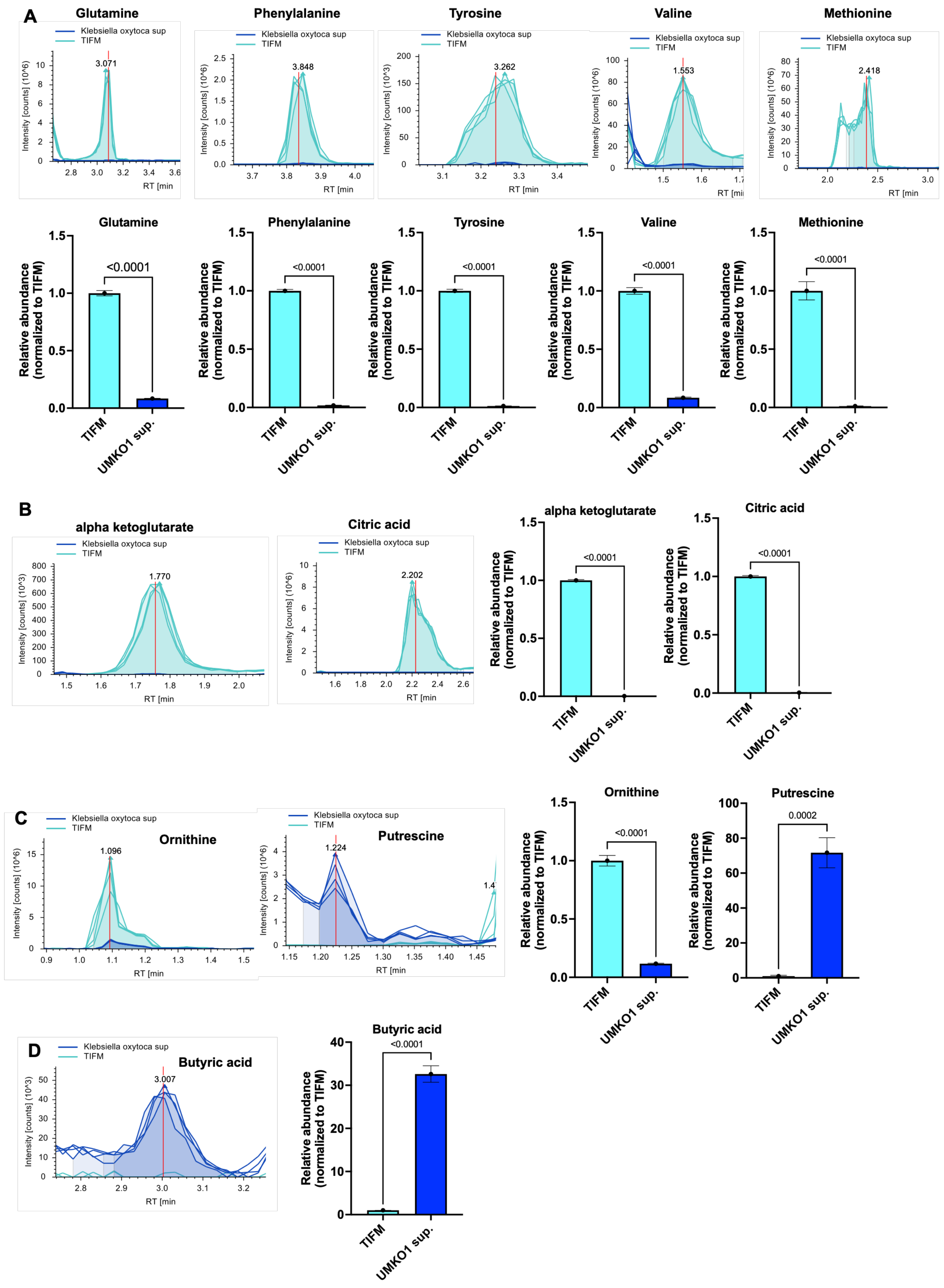


**Supplemental Figure 3: Metabolic features identified in the untargeted analysis.** Bacterial supernatant shows **(A)** depletion of several amino acids, as well as other key metabolites such as **(B)** alpha ketoglutarate and citric acid. **(C)** Ornithine, which is included in the TIFM was converted to putrescine by *Klebsiella oxytoca*. **(D)** The untargeted metabolomics was also able to pick-up short chain fatty acids such as butyric acid.

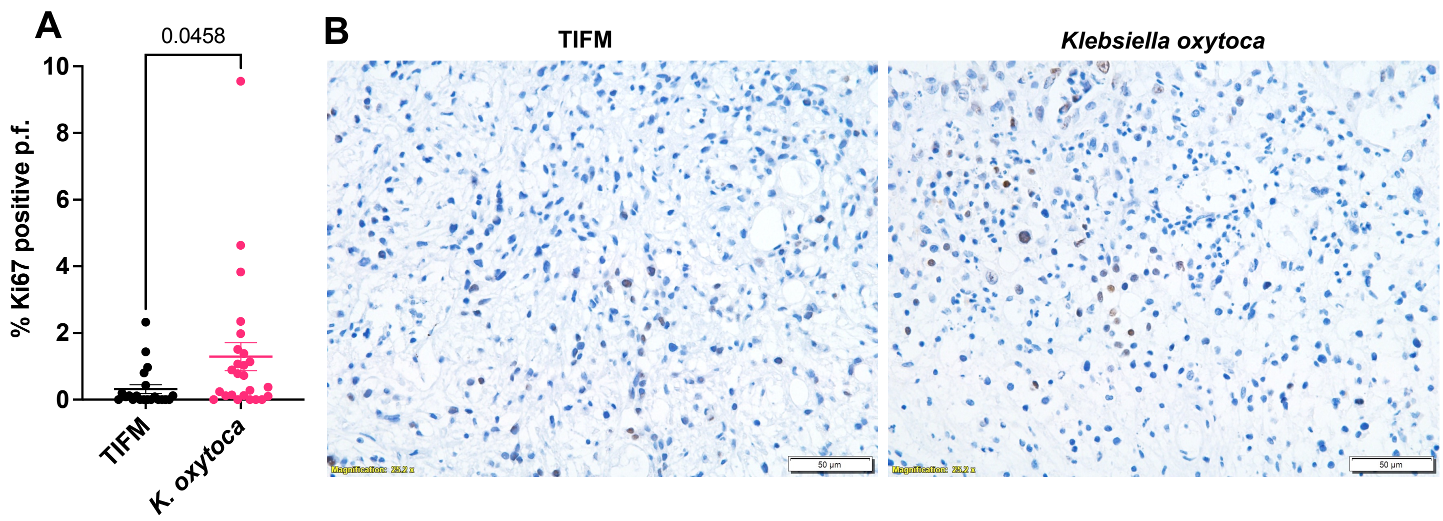


**Supplemental Figure 4: Tumor explants show an increase in Ki67 staining grown in the presence of *UMKO1* supernatant. (A)** Quantification of two tumor explants % Ki67 positive stained cells per field. **(B)** Representative images of the Ki67 stained tumor transplant system.
